# Transcription factor GLK2 regulates key steps of anthocyanin biosynthesis to antagonize photo-oxidative stress during greening of Arabidopsis seedlings

**DOI:** 10.1101/2023.03.10.532066

**Authors:** Xiyu Zeng, Luhuan Ye, Rui Zhang, Peng Wang

## Abstract

The greening of young seedling is light induced but vulnerable to photo-oxidative stress at the same time, due to the immature status of chloroplasts. Accumulation of anthocyanin is a protective response to high light, by absorbing excess energy and serving as antioxidant. In this work with Arabidopsis, we found that GARP family transcription factors GOLDEN2-LIKE 2 (GLK2), as a key regulator to chloroplast development, also plays an intensive role in regulating anthocyanin biosynthesis during seedling de-etiolation, especially under high light stress. We demonstrate that GLK2 positively regulates anthocyanin biosynthesis by directly activating the transcription of anthocyanin late biosynthetic genes (LBGs) as well as *TRANSPARENT TESTA GLABRA 1* (*TTG1*) gene, which encodes a key component of the regulatory MYB–bHLH–WDR (MBW) complex (which also activates LBGs). Our data further show that GLK2 and MBW each could activate the expression of *DFR* gene (key LBG) independently, via distinct promoter regions. We therefore propose a multifaceted involvement of GLK2 in anthocyanin biosynthesis as an important protective measure for developing chloroplasts, against excessive light exposure during seedling photomorphogenesis.

## Introduction

Light is a key signal that shapes and colors plant seedlings through a process called photomorphogenesis, which involves hypocotyl elongation, cotyledon opening, and the biosynthesis of phytohormones, chlorophyll, and anthocyanins. (Job & Datta, 2021; J. Li et al., 2020; X. Li et al., 2020; Nguyen et al., 2015). Plants often encounter light intensities that exceed their photosynthetic capacity (excess light). Excess light causes changes in the redox state of the photosynthetic electron transport chain and increases the production of reactive oxygen species (ROS). (Karpinski et al., 1997; Li et al., 2009; Xu et al., 2017).

A typical response to HL is the accumulation of Anthocyanins, a group of polyphenolic pigments that belong to the flavonoid group and found in plants as antioxidants as they are ROS scavengers (Agati et al., 2012; Yu et al., 2021). Anthocyanins can also absorb light to reduce the amount of energy reaching the photosynthetic apparatus (Agati et al., 2012; Page et al., 2012). Other abiotic stresses such as drought, salinity and extreme temperatures also lead to ROS generation, and anthocyanin accumulation under these conditions can confer tolerance through ROS scavenging (Agati et al., 2012; Nakabayashi et al., 2014).

Anthocyanin biosynthesis originates from a flavonoid synthetic pathway involving multiple enzymatic reactions. Anthocyanin early biosynthetic genes (EBGs) such as *CHALCONE SYNTHASE* (*CHS*), *CHALCONE ISOMERASE* (*CHI*), *FLAVANONE 3-HYDROXYLASE* (*F3H*), and *FLAVONOID 3’-HYDROXYLASE* (*F3’H*) are involved in the production of different types of flavonoids including anthocyanin; Anthocyanin late biosynthetic genes (LBGs) such as *DIHYDROFLAVONOL 4-REDUCTASE* (*DFR*), *LEUCOANTHOCYANIDIN OXYGENASE* (LDOX), *ANTHOCYANIDIN REDUCTASE* (*ANR*), and *UDP-GLUCOSE:FLAVONOID 3-O-GLUCOSYLTRANSFERASE* (*UF3GT*) are specific for anthocyanin biosynthesis only (Xu et al., 2015). Flavonoid biosynthesis is largely modulated at the transcriptional level, and the expression of anthocyanin LBGs is regulated by the MYB–bHLH–WDR (MBW) ternary transcriptional complex comprising 3 classes of regulatory proteins, including R2R3-MYB, basic helix-loop-helix (bHLH), and WD40-repeat proteins (WDR) (Li, 2014). A functional MBW complex is composed of TTG1 (the only WDR), one R2R3-MYB protein from PRODUCTION OF ANTHOCYANIN PIGMENTS (PAP1), PAP2, MYB113, or MYB114, and one bHLH protein from TT8, GLABROUS3 (GL3), or ENHANCER OF GLABRA3 (EGL3) (Baudry et al., 2004; Nesi, 2000; Shi & Xie, 2010; Tian & Wang, 2020; Xu et al., 2013; Zhou et al., 2020).

Anthocyanin accumulation in plants is modulated by light conditions (Chalker-Scott, 1999; Maier et al., 2013). The light regulation of anthocyanin biosynthesis involves several kinds of photoreceptors, such as PHYTOCHROMES A, B and E (PHYA/B/E) and CRYPTOCHROMES 1 and 2 (CRY1/2) (Chen et al., 2006; Warnasooriya et al., 2011). Downstream the photoreceptors, HY5 and PIF3 regulates the expression of anthocyanin biosynthesis genes through direct binding to the ACGT-containing elements (ACEs) and G-boxes in the promoters of both EBG and LBG, such as *CHS*, *CHI*, *F3H*, *F3’H*, *DFR*, and *LDOX* (Shin et al., 2007). These results suggest that anthocyanin biosynthesis is co-regulated and functionally active during photomorphogenesis.

GOLDEN2-LIKE transcription factors (GLKs) are key regulators of chloroplast development and are also involved in photomorphogenesis, photosynthesis, fruit development, and leaf senescence (Ahmad et al., 2019; Alem et al., 2022; Nguyen et al., 2014; Waters et al., 2008; Waters et al., 2009; Zhang et al., 2021). In Arabidopsis, a homologous pair of GLK1 and GLK2 redundantly regulates the expression of photosynthetic genes encoding chlorophyll biosynthesis, light-harvesting, and electron transport components, and only the *glk1 glk2* double mutants have pale-green photosynthetic tissues (Fitter et al., 2002; Waters et al., 2009). GLK gene pairs act redundantly to promote chloroplast development in Arabidopsis, however AtGLK2 has some functions that are independent of AtGLK1, like the AP1/CAL interaction (Fitter et al., 2002). Regarding the involvement of GLK1 in photomorphogenesis, a recent study showed that BRASSINOSTEROID INSENSITIVE2 (BIN2) phosphorylates GLK1 during seedling de-etiolation and promotes its stability to fine-tune chloroplast development (Zhang et al., 2021); TEOSINTE BRANCHED 1, CYCLOIDEA, and PROLIFERATING CELL FACTORS 15 (TCP15) interacts with GLK1 to control cotyledon opening (Alem et al., 2022). Nevertheless, the mechanisms and significance of GLK2 influence on photomorphogenesis remain largely unknown.

In previous studies in Arabidopsis, it was found that GLK1 acts upstream of MYBL2 to positively regulates sucrose-induced anthocyanin biosynthesis (Zhao et al., 2021), and GLK1 interacts with and enhances the transcriptional activation activities of MYB75, MYB90, and MYB113, resulting in increased expression of anthocyanin biosynthetic genes (Li et al., 2023). For GLK2, it was reported to positively regulates anthocyanin biosynthesis through HY5-mediated light signaling (Liu et al., 2021). However, the underlying molecular mechanism for GLK2 regulating anthocyanin accumulation remains unclear. Using de-etiolating Arabidopsis seedlings, we revealed that GLK2 acts downstream of HY5, to regulate both chloroplast development and hypocotyl elongation (unpublished work). In this study, we demonstrate that GLK2 positively regulate excessive light induced anthocyanin accumulation through direct transcriptional activation of the anthocyanin late biosynthetic genes as well as MBW regulatory component, helping to relief potential photo-oxidative stress encountered by Arabidopsis seedling during de-etiolation.

## Results

### GLK2 positively regulate anthocyanin accumulation against high light during seedling de-etiolation

To investigate the effect of GLK1 and GLK2 on anthocyanin biosynthesis during de-etiolation, we first analyzed anthocyanin content of normal light (NL, 100 μmol m^-2^ s^-1^) grown Arabidopsis seedlings. All the experimental materials were grown in darkness for 4 days, then transferred to continuous NL for 2 days in the growth chamber. We found that the anthocyanin content in *glk1* mutant have no different from the wild type (**Figure 1A, 1B**), while the anthocyanin content in *glk2* mutant was significantly lower than that in the wild type. The anthocyanin content in *glk1 glk2* double mutant decreased more significantly than that in *glk2* mutant. The *35S:GLK1* and *35S:GLK2* overexpression lines (in *glk1 glk2* double mutant background) both could compensate for the phenotype of decreased anthocyanin, but anthocyanin content increased more in *35S:GLK2* (**Figure 1A, 1B**).

**Figure 1.**
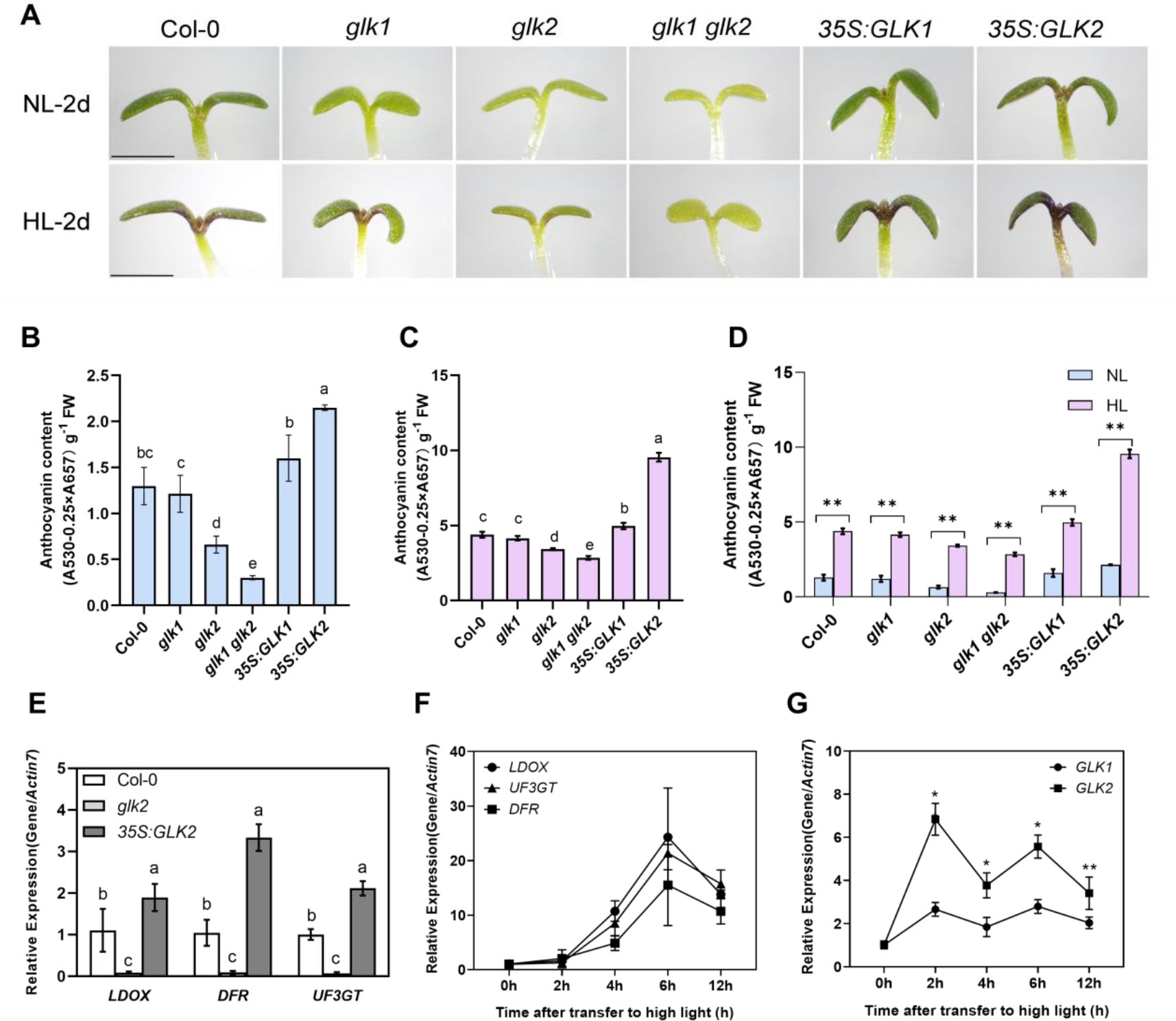
GLKs positively regulate anthocyanin accumulation during seedling de-etiolation in Arabidopsis **(A)** Phenotypes of wild-type (Col-0), *glk1*, *glk2*, double mutant (*glk1 glk2*), and double mutant lines overexpressing either *GLK1*(*35S:GLK1*) or *GLK2* (*35S:GLK2*). Seedlings germinated in darkness for 4 d were transferred to normal light (NL, 100 μmol m^-2^ s^-1^) and high light (HL, 300 μmol m^-2^ s^-1^) for 2 d in the growth chamber. Experiments were repeated three times and similar results were obtained. Scale represents 1 mm. **(B)** The anthocyanin content of NL grown seedlings were analyzed. **(C)** The anthocyanin content of HL grown seedlings were analyzed. **(D)** Comparison of the NL and HL effect on anthocyanin content. **(E)** Relative expression of anthocyanin late biosynthetic genes *DFR*, *LDOX*, and *UF3GT* in HL grown seedlings of Col-0, *glk2*, and *35S:GLK2*. **(F)** Relative expression of late anthocyanin biosynthetic genes in wild-type seedlings grown in HL for 0, 2, 4, 6, and 12 h. **(G)** Relative expression of *GLK* genes in wild-type seedlings grown in HL for 0, 2, 4, 6, and 12 h. The Actin7 gene was analyzed as an internal control, and the expression level of Col-0 was set to 1. Error bars represent mean±SD (n=3). Statistical analysis in (B), (C), and (E) was performed using ANOVA with Turkey’s HSD test; *P* < 0.05, different letters indicate statistically significant difference. Asterisks in (D) and (G) indicate significant differences with \**P* < 0.05 and \*\**P* < 0.01 (Student’s *t*-test), respectively.

High light induces anthocyanin accumulation in plants against excess light and ROS (Agati et al., 2012; Page et al., 2012). We next transferred etiolated seedlings to high light (HL, 300 μmol m^-2^ s^-1^), which induced more anthocyanin accumulation in all materials (**Figure 1D**). We find that anthocyanin content in *glk1* mutant remained no difference from the wild type. Although lower than wild type in anthocyanin content, the relative difference between *glk2*, *glk1glk2* and the wild type was smaller. In contrast, the anthocyanin content in *35S:GLK2*, but not *35S:GLK1* overexpression line, showed dramatic accumulation compared with all other materials (**Figure 1A, 1C**). Similar trend was found in materials after 24 hours de-etiolation under HL (**Supplemental Figure 1**). The difference between *35S:GLK2* and *35S:GLK1* in anthocyanin accumulation especially under HL, indicate that GLK2 may be more directly regulating anthocyanin biosynthesis in Arabidopsis. We therefore focus on GLK2 for the following studies.

### High light induces *GLK2* early expression and *GLK2* deficiency retards seedling greening and anthocyanin accumulation

We analyzed the expression of anthocyanin late biosynthetic genes (LBGs) in wild-type (Col-0), *glk2* mutant, and *35S:GLK2* overexpression line after seedling de-etiolation under continuous HL for 2 days. RT-qPCR analysis showed that the expression of anthocyanin LBGs (*DFR, LDOX* and *UF3GT*) were all significantly higher in *35S:GLK2* overexpression line, and were all significantly lower in *glk2* mutant, than those in the wild type (**Figure 1E**). This result further indicates that GLK2 may directly activates anthocyanin late biosynthetic genes.

During early stage seedling de-etiolation, the expression levels of LBGs changed drastically between 2-6 h of HL illumination (**Figure 1F**). The expression levels of *GLK1* and *GLK2* genes both increased along the same time course, but the gene expression of *GLK2* is higher than *GLK1* at all time points. Interestingly, HL treatment induced a faster increase of *GLK* gene expression than NL treatment, especially that the expression of *GLK2* quickly reached a remarkably higher level compared with *GLK1* at as early as 2 h upon HL illumination (**Figure 1G**, **Supplemental Figure 2**). These results suggest that compared with *GLK1*, *GLK2* exhibits faster response to HL, indicating its protective role related to anthocyanin biosynthesis during seedling de-etiolation in Arabidopsis.

For further assessment of GLK2 function during early stage of de-etiolation, we analyzed the phenotypes of seedling greening by using a whole set of *glk* mutants and overexpression lines under continuous normal light condition. Retarded seedling greening was found in *glk2* mutant starting from 12 hours of de-etiolation, precocious seedling greening was found in *35S:GLK2* overexpression line starting from 4 hours of de-etiolation, and anthocyanin accumulation was visible in *35S:GLK2* at 24 hours (**Supplemental Figure 3**). We then analysed the wild type, *glk2* mutant, and *35S:GLK2* overexpression line after 12 h of seedling de-etiolation under NL and HL. Results showed that HL treatment further retarded seedling greening of *glk2*, and anthocyanin accumulation in *35S:GLK2* was brought forward to be visible at 12 hours, while anthocyanin accumulation in *glk2* mutant was significantly lower (**Figures 2A, 2B**). We thus suspect that the paler cotyledon in *glk2* was at least partially attributable to high light induced photoxidative stress, apart from the deficient in chlorophyll biosynthesis (Fitter et al., 2002; Waters et al., 2009).

**Figure 2.**
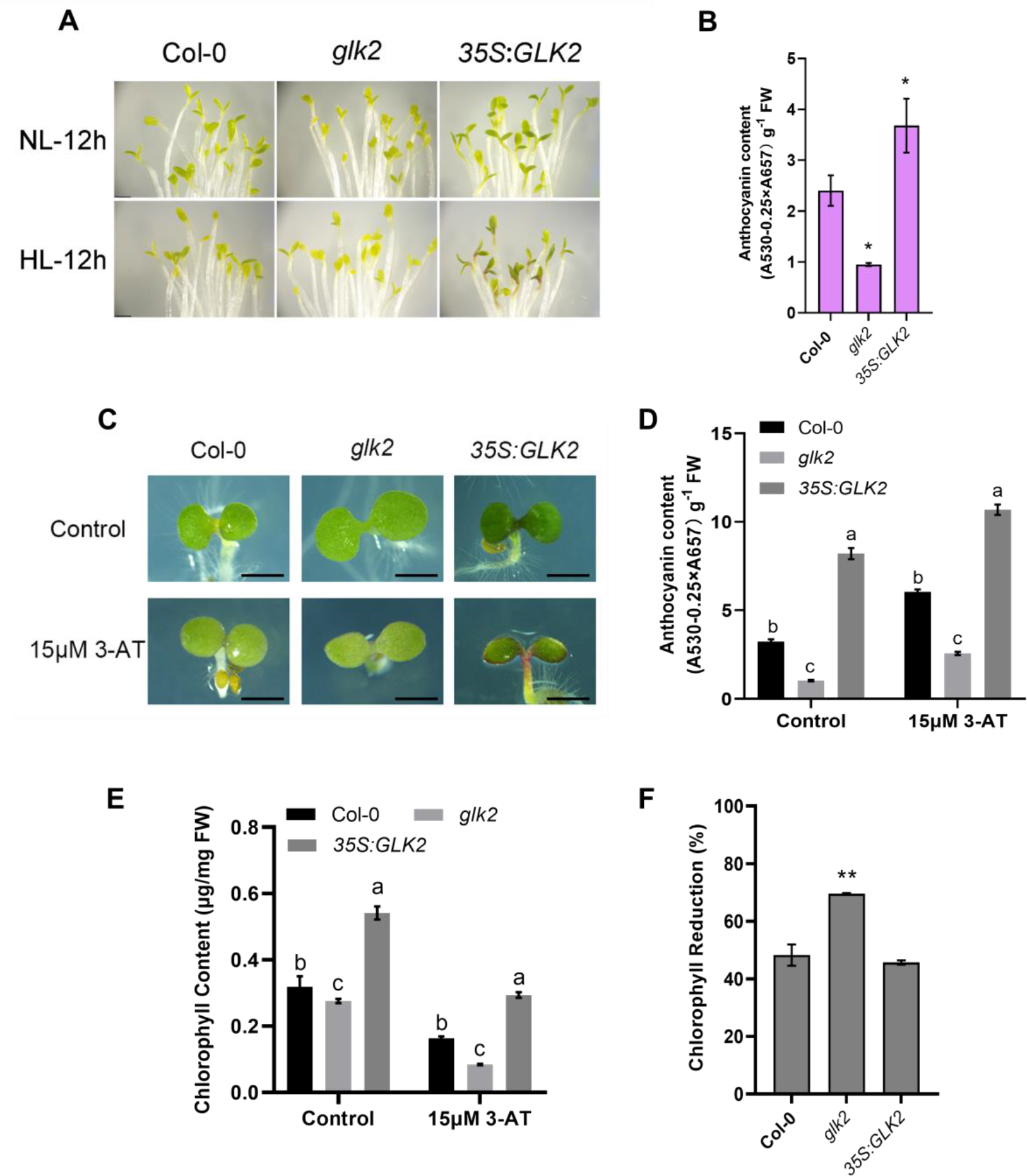
Loss of AtGLK2 reduces anthocyanin accumulation and is responsible for photo-oxidative chlorophyll reduction **(A)** Phenotypes of wild-type (Col-0) , *glk2* and double mutant lines overexpressing *AtGLK2* (*35S:GLK2*) seedlings after germination in darkness for 4 d and transferred to high light (HL, 300 μmol m^-2^ s^-1^) for 12 h in the growth chamber. **(B)** The anthocyanin content of the 12 h HL grown seedlings were analyzed. **(C)** Phenotypes of Col-0, *glk2*, and *35S:GLK2* seedlings germinated and grown in normal light (NL, 100 μmol m^-2^ s^-1^) for 4 d, under control condition (1/2 MS) and 15μM 3-AT treatment. **(D, E)** The anthocyanin and chlorophyll contents of the samples from (C). **(F)** The extent of Chl reduction of the 3-AT treated relative to the control samples. Phenotypic experiments were repeated three times and similar results were obtained. Scale bar represents 1 mm. Error bars in (B) represent the mean±SD (n=3), and in (D), (E), and (F) represent the means±SD (n=5). Asterisks in (B) and (F) indicate significant differences (\**P* < 0.05; \*\**P* < 0.01). Statistical analysis in (D) and (E) were performed using ANOVA with Turkey’s HSD test; *P* < 0.05, different letters indicate statistically significant difference.

### De-etiolating GLK2 deficient mutants are sensitive to ROS induced chlorophyll reduction

Previous study have revealed that the ROS-scavenging role of anthocyanin prevents chlorophyll loss and helps in maintaining photosynthetic capacity, while anthocyanin deficient mutants accumulate more ROS (Xu & Rothstein, 2018). ROS was produced in chloroplasts via both photosystems due to the excess photons trapped in photosystem (PSII) and the electrons transferred to molecular oxygen through photosystem I (PSI) (Khorobrykh et al., 2015; Triantaphylidès & Havaux, 2009). Electron leakage on the PSII electron acceptor side produces O_2_^-^ , which undergoes dismutation to H_2_O_2_ by superoxide dismutase (SOD) (Mhamdi & Van Breusegem, 2018). H_2_O_2_ is then converted to water and dioxygen by peroxidase and catalase (CAT). We used 3-amino-1,2,4-triazole (3-AT, a CAT inhibitor, thus triggering accumulation of H_2_O_2_) to induce oxidative stress. After germinating seeds of wild-type (Col-0), *glk2*, and *35S:GLK2* on 1/2 MS with 15 μM 3-AT for 4 d, we observed that Col-0 and *glk2* mutant cotyledons turned yellow compared with the control, whereas *35S:GLK2* cotyledon stayed green with dramatic accumulation of anthocyanin (**Figure 2C**). 3-AT treatment greatly induced the anthocyanin accumulation and reduced the chlorophyll accumulation in all experimental materials, and notably, *glk2* mutant exhibited a larger extent of chlorophyll reduction compared with the wild-type (**Figures 2D, 2E, 2F)**. These results indicate that anthocyanin-defective *glk2* mutants are more sensitive, whereas anthocyanin-over-accumulated *35:GLK2* overexpression lines are more tolerant to 3-AT induced oxidative stress than the wild-type.

In mature Arabidopsis plants (5-week-old), anthocyanin accumulates significantly lower in *glk1 glk2* double mutants, but significantly higher in the *35S:GLK2* overexpression line, than that of WT in the petiole (**Supplemental Figure 4A, 4B**). We also treated 5-week-old Arabidopsis under high light (500 μmol m^-2^ s^-1^) for 2 d, and similar trends were found although anthocyanin accumulation increased in all materials (**Supplemental Figure 4C**), which is consistent with the results from high light treated etiolating seedlings. H_2_O_2_ content appeared similar in *35S:GLK2* overexpression line and wild-type, while lower in the double mutant (**Supplemental Figure 4D**), indicating more complicated ROS generation and scavenging systems are operating in these plants at mature stage. If assuming more ROS generation in *35S:GLK2* due to higher density of photosynthetic apparatus, the abundance of anthocyanin might be responsible to stable ROS accumulation relative to the wild-type.

### GLK2 transcription factor directly binds and activates the promoters of anthocyanin LBGs

A core binding motif of RGATTYY conserved to GOLDEN2-LIKE (GLK) transcription factors was identified from previous study (Tu et al., 2022). In this work we mainly use CCAATC and GATTCT sequences for binding tests, as they have been studied before in Arabidopsis (Ahmad et al., 2019; Waters et al., 2009; Zhang et al., 2021). Promoter analysis of anthocyanin LBGs (around 2 kb upstream of the translational start site) revealed several CCAATC and GATTCT sequences (**Figure 3A**). Electrophoretic mobility shift assays (EMSAs) were performed to determine whether GLK2 binds to these elements. The result shows, the purified GLK2 recombinant protein bound to the promoters of *DFR, LDOX and UF3GT*, competitive probes weakened such binding, while the binding activity wasn’t diminished when a mutant competitive probe was used (**Figures 3B, 3C**). While obtaining the above *in vitro* results, to further confirm that GLK2 binds *in vivo* to the promoters of anthocyanin LBGs, chromatin immunoprecipitation (ChIP) assays were performed using a GLK2 antibody to immune-precipitate GLK2-DNA complexes from *glk1 glk2* mutants and 35S:GLK2 overexpression line. Enriched DNA sequences were amplified by RT-qPCR using primer pairs covering the promoter regions of anthocyanin LBGs, to confirm that the native GLK2 indeed bound to the promoters of anthocyanin LBGs. Results demonstrated that the D and G regions from *DFR* promoter, the B region from *LDOX* promoter, and the B region from UF3GT promoter were most strongly bound by GLK2 (**Figure 3D**). We then performed transient transcription assays to examine whether these sequences were required for transcriptional activation of anthocyanin LBGs by GLK2. Dual-luciferase reporter plasmids containing the Firefly luciferase gene driven by *DFR, LDOX, or UFGT* promoters, and the Renilla luciferase gene driven by the constitutive 35S promoter were used in the assays (**Figure 3E**). The effector and reporter were co-expressed in Arabidopsis protoplasts. Compared with the control (empty effector), the GLK2 effector construct significantly promoted the luciferase reporter reactions, indicating that GLK2 directly activates the transcription of anthocyanin LBGs (**Figure 3F**).

**Figure 3.**
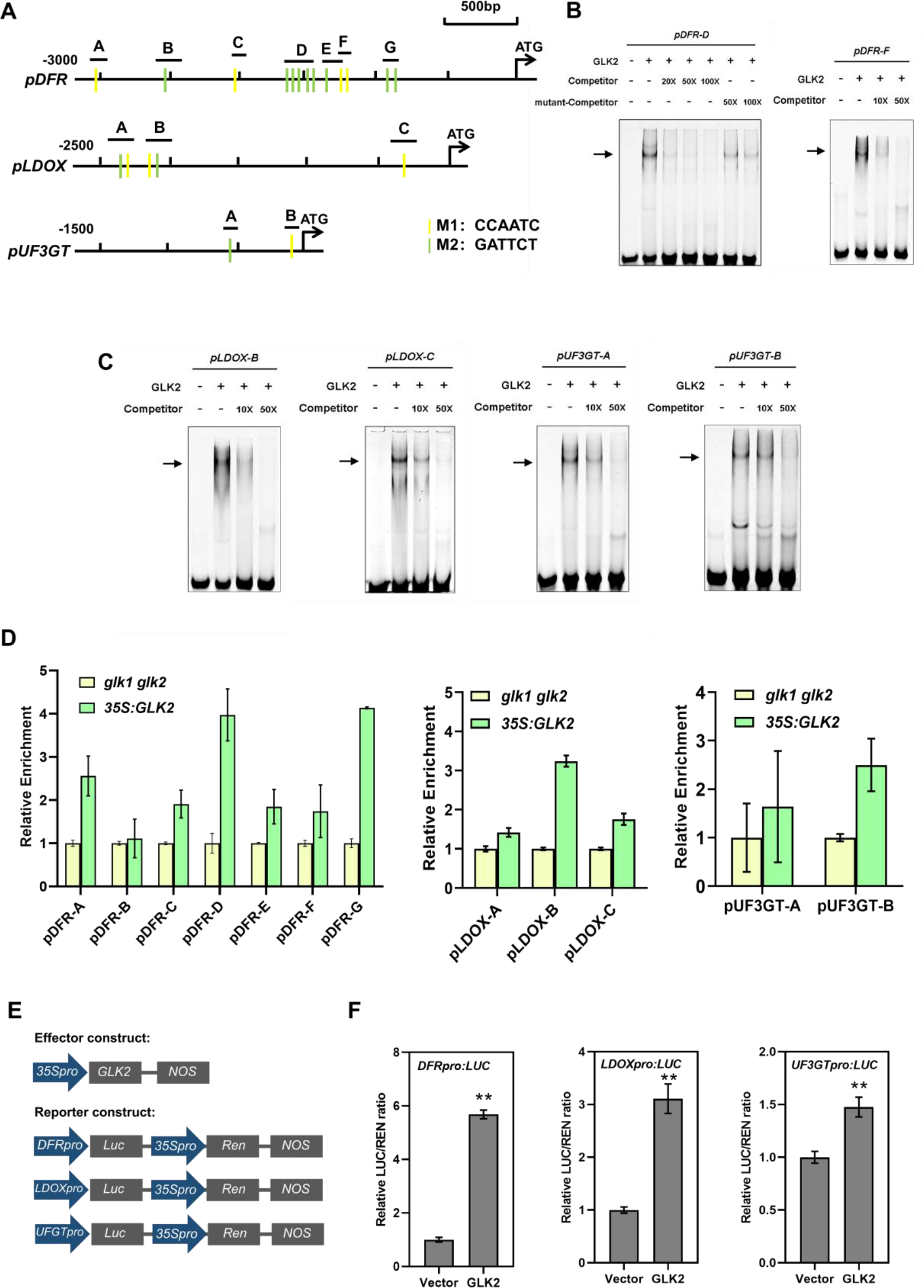
GLK2 binds to the promoters and activate the expression of anthocyanin late biosynthetic *DFR* , *LDOX*, and *UF3GT* genes **(A)** Illustration of the *DFR*, *LDOX*, and *UF3GT* promoter regions showing the presence of putative GLK binding motifs. Yellow strings indicate CCAATC (M1) elements and green strings indicate GATTCG (M2) elements in the promoters. The letters indicate primer pairs used for ChIP-qPCR or EMSA assays. **(B, C)** EMSA assays showing that GLK2 protein binds to the *DFR*, *LDOX* and *UF3GT* promoters *in vitro*. The probes (59bp, DFR-probeD, -1541bp ∼ -1483bp; 74bp, DFR-probeF, -1295bp ∼ -1222bp; 48 bp, LDOX-probeB, -2191bp∼-2122bp; 75bp,LDOX-probeC, -357bp∼-301bp; 56bp, LUF3GT-probeA, -611bp∼-555bp; 56bp, LUF3GT-probeB, -158bp∼-102bp; 56bp.) were labeled with Cy5. Unlabeled probe was added as a competitor. pDFR-p1m is a mutant probe (CCAATC mutated to AAAAAA) which could not be bound. Arrows indicate that GLK2 binds to the probes. **(D)** ChIP-qPCR results showing that GLK2 binds to the promoters of *DFR*, *LDOX* and *UF3GT in vivo*. Two-week-old *glk1 glk2* double mutant and double mutant lines overexpressing *AtGLK2* (*35S:GLK2*) transgenic plants were grown in half-strength MS medium in the growth chamber before harvesting samples. Chromatin fragments (∼500 bp) were immunoprecipitated by GLK2 antibody coupled Dynabeads (IP). The precipitated DNA was analyzed by RT-qPCR using primer pairs based on the promoters of *DFR*, *LDOX* and *UF3GT*. The level of binding was calculated as the ratio between IP and input, normalized to that of Actin7 (internal control), and the binding level in *glk1 glk2* double mutant was set to 1. Error bars represent mean±SD (n=3). **(E)** Structure of the *DFR*, *LDOX*, and *UF3GT* promoter-driven dual-luciferase (LUC) reporter constructs and 35S promoter-driven GLK2 effector construct. For the reporter constructs, *DFR* promoter (-1701 bp to -1222 bp), *LDOX* promoter (-2478 bp to -1 bp), and *UF3GT* promoter (-344 bp to -30 bp) are used. **(F)** Dual-luciferase assays showing GLK2 promote the expression of *DFR*, *LDOX* and *UF3GT* genes. Arabidopsis protoplasts isolated from Col-0 were co-transfected with reporter DNA and effector DNA. After transfection, the protoplasts were kept in low light for 16 h. LUC activity was normalized to REN activity. The ratio of LUC/REN in empty effector vector were set to 1. Error bars represent mean ± SD (n=3). Asterisks indicate a significant difference compared with control vector (\**P*<0.05, \*\**P*<0.01, Student’s *t*-test).

### GLK2 positively regulates MBW complex member genes and directly activates *TTG1*

The expression of anthocyanin LBGs is regulated by MYB–bHLH–WDR (MBW) complex, which includes members of R2R3-MYB, bHLH, and WDR transcription factors (**Figure 4A**) (Li, 2014). After confirming direct regulation of anthocyanin LBGs by GLK2, we moved on to test whether GLK2 affect MBW expression. Using 2d HL treated de-etiolating seedlings, we found that the expression levels of R2R3-MYB transcription factors *PAP1* and *PAP2*, bHLH transcription factors *TT8*, *GL3 and EGL3* were significantly lower in *glk2* mutant, and significantly higher in *35S:GLK2* overexpression line than those in the wild type (**Supplemental Figure 5**). The result indicates that GLK2 might promote MBW complex function during seedling de-etiolation in Arabidopsis.

**Figure 4.**
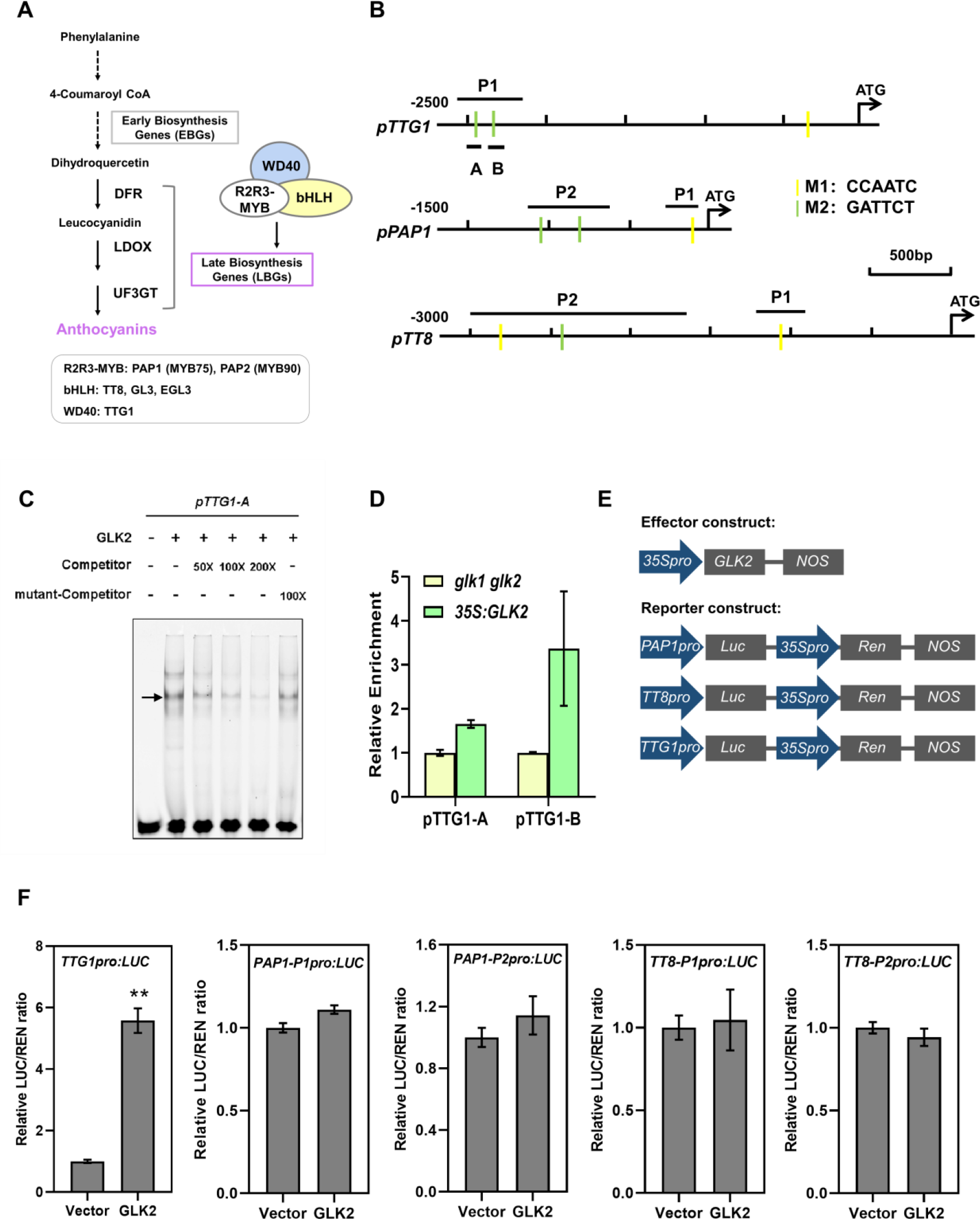
GLK2 binds to the promoter and activates the gene expression of MBW component TTG1 **(A)** Schematics showing the regulatory involvement of MBW complex (R2R3-MYB, bHLH, WD40) in anthocyanin biosynthetic pathway. **(B)** Illustration of the *TTG1*, *PAP1*, and *TT8* promoter regions showing the presence of putative GLK binding motifs. Yellow strings indicate CCAATC (M1) elements and green strings indicate GATTCG (M2) elements in the promoters. The letters indicate primer pairs used for EMSA assay, ChIP-qPCR, or dual-luciferase assays. **(C)** EMSA assays showing that GLK2 protein binds to the *TTG1* promoter *in vitro*. The TTG1 probe (45bp; -2514bp ∼ -2470bp) were labeled with Cy5. Unlabeled probe was added as a competitor. pTTG1m is a mutant probe (GATTCT mutated to AAAAAA) which could not be bound. The arrow indicates that GLK2 binds to the probes. **(D)** ChIP-qPCR results showing that GLK2 binds to the promoter of *TTG1 in vivo*. The experiment was performed same as in Figure 3D. Error bars represent mean±SD (n=3). **(E)** Structure of the *PAP1*, *TT8*, and *TTG1* promoter-driven dual-luciferase (LUC) reporter constructs and 35S promoter-driven GLK2 effector construct. For the reporter constructs, *PAP1* promoters (-258 bp to -53 bp), *TT8* promoters (-1398 bp to -1096 bp), and *TTG1* promoters (-2578 bp to -2155 bp) are used. **(F)** Dual-luciferase assays showing GLK2 promote the expression of *TTG1* but not *PAP1* and *TT8* genes. The assays were performed same as in Figure 3F. Error bars represent mean ± SD (n=3). Asterisks indicate a significant difference compared with control vector (\*\**P*<0.01, Student’s *t*-test).

We further analyzed the promoter regions of MBW complex member genes (up to 3kb upstream of the translational start site), and several CCAATC and GATTCT sequences for GLK binding motif were found (**Figure 4B**). To examine whether GLK2 directly regulates the gene expression of MBW complex members, we performed transient transcription assays using dual-luciferase reporter system constructed for *PAP1*, *TT8*, and *TTG1* genes (**Figure 4E**). Results showed that GLK2 significantly promoted the luciferase reporter reactions for *TTG1* but not for *PAP1* and *TT8* (**Figure 4F**), indicating that GLK2 directly promotes the gene expression of WDR transcription factor TTG1. On the basis of this, the physical binding of GLK2 to *TTG1* gene promoter was proofed by EMSA assay *in vitro* (**Figure 4C**), and ChIP-qPCR assay *in vivo* (**Figure 4D**).

### GLK2 activates *DFR* gene expression independent from MBW complex via distinct promoter region

GLK transcription factors are members of the GARP family of Myb transcription factors (Hosoda et al., 2002). The sequence of GARP DNA binding domain is similar to the TEA-binding domain and to the MYB-like domain in MYB-related proteins (Baranowskij et al., 1994). Interactions between bHLH- and MYB-type plant proteins have been widely documented (Nesi, 2000; Wen et al., 2018). Therefore, we examined whether GLKs interact with the bHLH protein TT8. Tagged GLK1, GLK2, and TT8 proteins were expressed and purified from *E. coli.*, and pulldown assay showed that GLKs physically interact with TT8 *in vitro* (**Figure 5A**). The GLK2 and TT8 interaction was further proved by strong luciferase activity in tobacco leaves co-infiltrated with GLK2-nLUC and cLUC-TT8 plasmids (**Figure 5B**).

**Figure 5.**
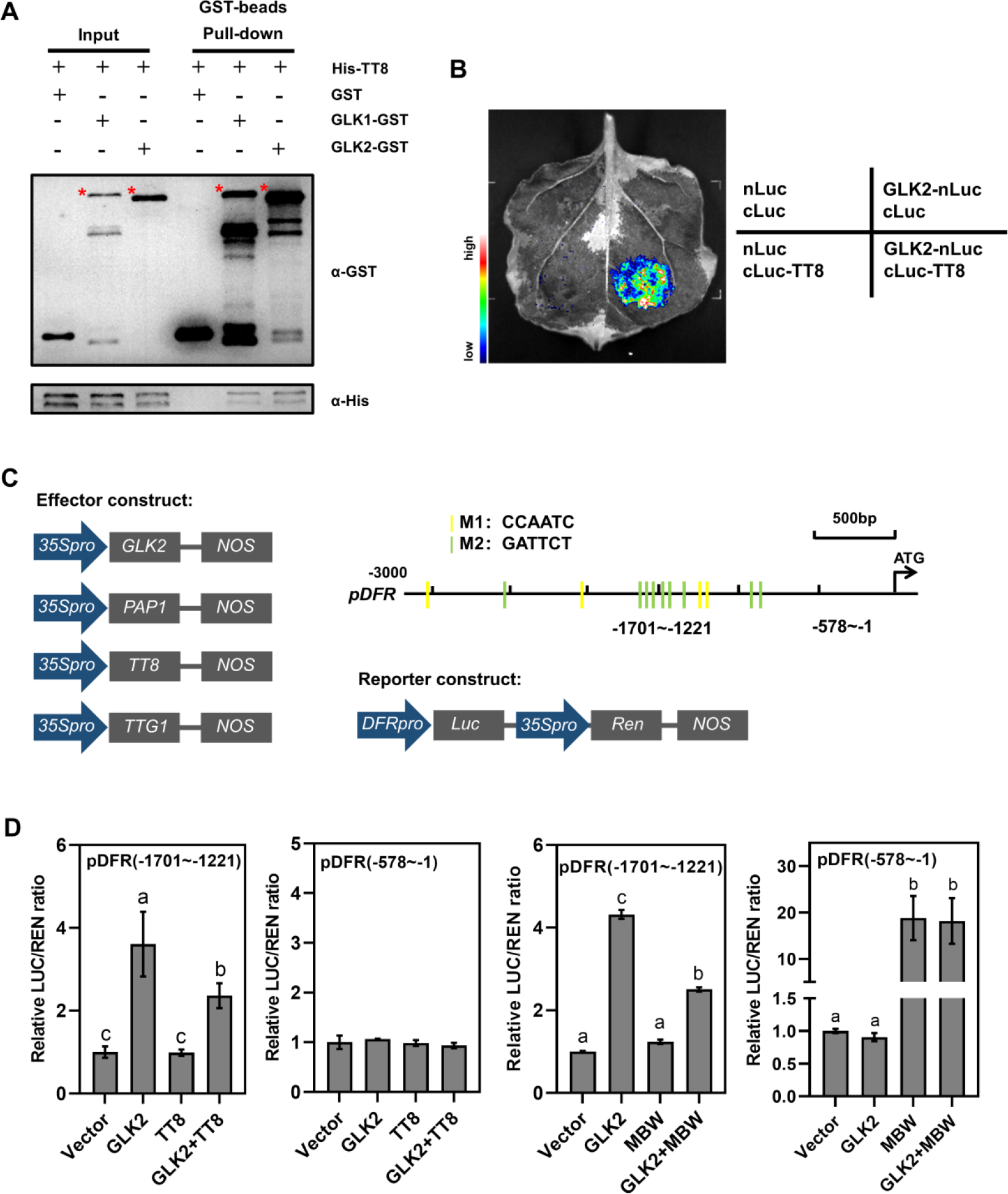
GLK2 and MBW complex activate *DFR* gene via distinct promoter regions **(A)** *In vitro* pulldown assays showing the interaction of GLK1 and GLK2 with TT8. GST-GLK protein or GST protein were used to pull down His-TT8 protein using GST beads. Anti-GST and anti-His antibodies were used for immunoblot analysis. “-” and “+” indicate the absence and presence of corresponding proteins. **(B)** Bimolecular luminescence complementation assay showing GLK2 interacts with TT8. nLUC and cLUC served as negative controls. **(C)** Structure of the *DFR* promoter-driven dual-luciferase (LUC) reporter construct and 35S promoter-driven GLK2, PAP1, TT8, and TTG1 effector constructs. For the reporter constructs, *DFR* promoter regions of -578 bp to -1 bp and -1701 bp to -1222 bp are used. Also illustrated the *DFR* promoter regions showing the presence of putative GLK binding motifs, and the indicated regions of -1701∼-1221 bp and -578∼-1 bp were used for dual-luciferase assays. **(D)** Dual-luciferase assays showing GLK2 and MBW complex each could active the expression of *DFR* gene independently, via distinct promoter regions of -1701∼-1221 bp and -578∼-1 bp respectively. The assays were performed similar as in Figure 3F, with the effector DNA “MBW” composed of “PAP1 + TT8 + TTG1”, and “GLK2 + MBW” composed of “GLK2 + PAP1 + TT8 + TTG1”. Error bars represent mean±SD (n=3). Letters “a” to “c” indicate statistically significant difference, as determined by one-way ANOVA, followed by Tukey’s HSD test (P < 0.05).

**Figure 6.**
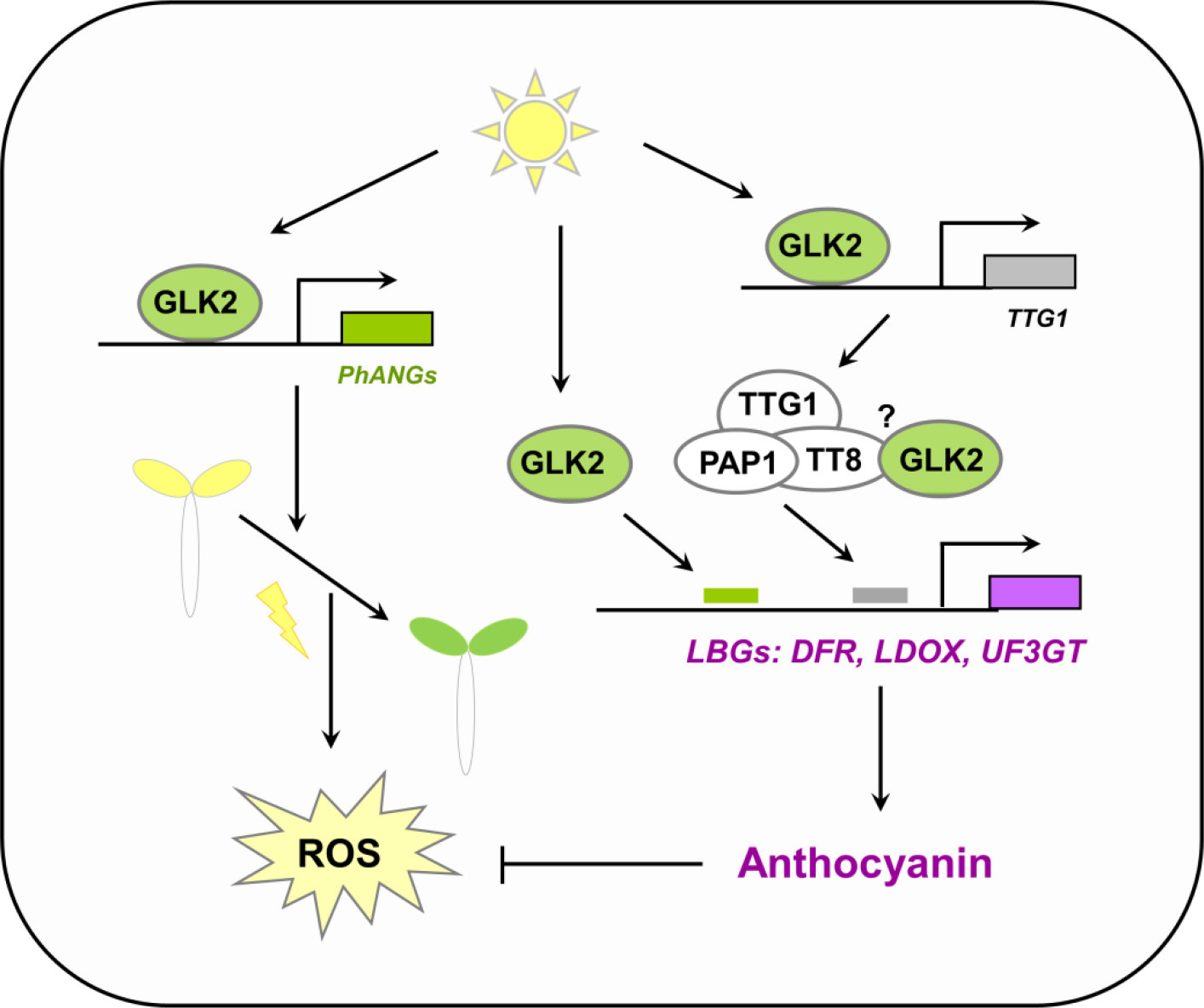
A model of GLK2 transcription factor promotes anthocyanin biosynthesis to antagonize photo-oxidative damage during seedling greening. During de-etiolation, GLK2 promotes chlorophyll biosynthesis and chloroplast development, simultaneously GLK2 promotes anthocyanin biosynthesis through directly activates anthocyanin late biosynthetic genes (LBGs). It also activates the gene expression of TTG1 and may interact with TT8, both are important components of MBW complex, which transcriptionally regulates LBGs. Our data shows that GLK2 and MBW each could activate the expression of *DFR* gene independently, via distinct promoter regions. Excess light causes increased production of reactive oxygen species (ROS) in greening seedlings, while the accumulation of anthocyanins acts as antioxidant or ROS scavengers. We therefore propose a multifaceted involvement of GLK2 in anthocyanin biosynthesis as an important protective measure for developing chloroplasts, against excessive light exposure during seedling photomorphogenesis. Arrows and bars represent positive and negative regulation, respectively.

To test whether GLK2 and TT8 interaction affects their regulations to target genes, we continue to use dual-luciferase reporter system, but with single or combination of effector constructs for comparison. In addition, DFR promoter-driven reporter constructs with promoter regions of -578 bp to -1 bp (this region includes TT8 binding site but without GLK binding motif), and -1701 bp to -1221 bp (this region includes several GLK binding motifs but without TT8 binding site) were used for activation assays (**Figure 5C**). We first transiently expressed the DFRpro:LUC (-578∼-1) reporter in wild-type protoplasts together with either 35S:GLK2 or 35S:TT8 or combination of both effectors, and the result showed the expression of DFRpro:LUC (-578∼-1) wasn’t activated by any of the added effectors. Then we transiently expressed the other DFRpro:LUC (-1701∼-1221) reporter together with either 35S:GLK2 or 35S:TT8 or both. The result showed the expression of DFRpro:LUC (-1701∼-1221) was significantly activated by GLK2 but not TT8, and the combination of both resulted in lower expression than that caused by GLK2 effector (**Figure 5D**).

We further transiently expressed the different DFRpro:LUC reporters together with either 35S:GLK2 or MBW complex (including 35S:PAP1, 35S:TT8 and 35S:TTG1) or both. When DFRpro:LUC (-578∼-1) reporter was used, the expression of it wasn’t activated by GLK2 but distinctly activated by MBW complex. After the addition of GLK2, the expression didn’t change. On the contrary, when DFRpro:LUC (-1701∼-1221) reporter was used, the expression of it was activate by GLK2 but not MBW complex. After the addition of MBW complex, the expression was lower (**Figure 5D**). Moreover, we also transiently expressed the DFRpro:LUC (-1701∼-1221) reporter together with either 35S:GLK2 or 35S:PAP1 or 35S:TTG1, or 35S:GLK2 plus 35S:PAP1, or 35S:GLK2 plus 35S:TTG1 (**Supplemental Figure 5**). Result showed the expression of DFRpro:LUC (-1701∼-1221) was activate by GLK2 but neither PAP1 nor TTG1. After the addition of PAP1 or TTG1 to GLK2, the expression of the reporter reduced. Taken together, these results suggest that GLK2 and MBW complex each could active the expression of *DFR* gene independently, via binding to distinct promoter regions.

## Discussion

In this work, we focus to dissect the molecular mechanism behind the intensive role of GLK2 in promoting anthocyanin accumulation, which provide protection against high light stress during seedling de-etiolation in Arabidopsis. We propose that the mechanism and the extent of GLK2 regulating anthocyanin biosynthesis likely differ from that of GLK1, which was reported recently (Li et al., 2023).

During Arabidopsis seedling de-etiolation, high light effectively increased anthocyanin accumulation in wild-type, whereas the effects were impaired in *glk2* mutant but enhanced in *GLK2* overexpression line, indicating that GLK2 is a positive regulator for high light induced anthocyanin accumulation. Consistently, transcript levels of anthocyanin late biosynthetic genes (LBGs) and regulatory pathway, including *DFR*, *LDOX*, and *UF3GT*, as well as *PAP1*, *PAP2*, *TT8*, *EL3*, *EGL3*, and *TTG1* were significantly decreased in *glk2* mutant but increased in *35S:GLK2* compared with the wild-type, indicating that GLK2 is transcriptional activator of these genes. Transient expression assays revealed that GLK2 could activate the promoters of *DFR*, *LDOX*, *UF3GT*, and *TTG1* in Arabidopsis protoplasts. Moreover, chromatin immunoprecipitation-quantitative PCR (ChIP-qPCR) assay and electrophoretic mobility shift assay (EMSA) showed that GLK2 directly binds to GLK core binding motif RGATTYY (CCAATC or GATTCT) present in the promoters of *DFR*, *LDOX*, *UF3GT* and *TTG1*. Taken together, these results suggest that GLK2 positively regulate high light induced anthocyanin accumulation through transcriptional activation of the anthocyanin late biosynthetic genes and regulatory pathway.

We further answered the question that whether GLK2 also indirectly regulates anthocyanin biosynthesis by activating other transcription factors and together with them to co-regulate downstream anthocyanin biosynthetic genes. Previous study noticed that gene expression of MBW components PAP1 and TT8 was positively regulated by GLK2, and speculated that they might be co-factors of GLK2 in Arabidopsis (Liu et al., 2021). In our work, the gene expression of all the tested MBW complex members, including PAP1, PAP2, TT8, GL3, EGL3 was observed to be significantly higher in *35S:GLK2* overexpression lines than those in the wild type (**Supplemental Figure 4**), but only the transcription of *TTG1* gene was further proved to be directly activated by GLK2 (**Figure 4**). The expression levels of *PAP1*, *PAP2*, and *TT8* were lower in *glk2* mutant than those in the wild type, indicating the negative effects of GLK deficiency on them (**Supplemental Figure 4**). It is possible that they are indirectly affected by GLK2, or GLK2 interacts with other proteins to be able to activate their transcriptions. The WDR protein TTG1 functions as a scaffold for DNA-bound MYB and bHLH proteins to interact to generate the MBW complex, activating the expression of anthocyanin-biosynthetic genes (Zhang & Schrader, 2017). There is no DFR expressed in *ttg1* mutants (Shirley et al., 1995), while the TTG1-dependent MBW complexes are able to bind directly to the promoters of *TTG2*, *TT8*, *F3’H*, *DFR*, and *LDOX* genes to regulate their expression (Lee et al., 2019). The regulation of *TTG1* by GLK2 suggests an indirect GLK2 regulatory pathway on anthocyanin LBGs.

As the direct transcriptional regulation of PAP1, PAP2, TT8, GL3, and EGL3 by GLK2 is currently uncertain, we then tested the protein interaction relationships between GLK2 and these MBW members. Among the 6 (including TTG1) tested by BiLC assay, only TT8 was shown to interact with GLK2, and further confirmed by protein pull-down assay (**Figure 5A, 5B**). Although GLK2 can physically interact with TT8 *in vitro*, GLK2 wasn’t affecting the transcriptional activation activities of TT8 or MBW complex (the combination of PAP1, TT8 and TTG1) on *DFR* gene expression, and *vice versa*, so that the significance of this interaction remain to be discovered. In fact, GLK2 and MBW complex each could independently activate *DFR* promoter, although MBW does with higher extent (**Figures 5C, 5D**).

GLKs are master regulators involved in chlorophyll biosynthesis and chloroplast development, but to what extent they participate in photomorphogenesis is not clear. HY5 is the regulatory hub in light signaling and photomorphogenesis. One of our parallel work recently reveals that GLKs act downstream of HY5, to regulate both chloroplast development and hypocotyl elongation in Arabidopsis, which are two of the most important processes during photomorphogenesis, adding to the recently published work showing that GLK1 controls cotyledon opening (Alem et al., 2022). Together with our finding of direct regulation of GLK2 on anthocyanin biosynthesis, all these facts and clues suggest an important regulatory cascade from HY5, GLK, to multifaceted photomorphogenesis processes. To summarize this study, the high light induced GLK2 early expression, effects of GLK2 deficient on seedling greening and anthocyanin accumulation, and the GLK2 participation in regulation of both anthocyanin LBGs and MBW complex, suggest a further prominent role of GLK2 in Arabidopsis seedling photomorphogenesis.

## Materials and methods

### Plant materials and growth conditions

*Arabidopsis thaliana* ecotype Columbia (Col-0) was used in all experiments. Mutants of *glk1*, *glk2* and *glk1 glk2* (Fitter et al., 2002), and overexpression lines of *35S:GLK1* and *35S:GLK2* in double mutant background (Waters et al., 2008) are described previously. The surface-sterilized seeds were treated at 4°C in the dark for 3 days, and germinated on half-strength Murashige and Skoog (MS) medium (pH 5.8) containing 0.8% agar and 1% sucrose at 22°C in darkness. Four days after germination, the etiolated seedlings were transferred to continuous normal light (NL, 100 µmol m^−2^ s^−1^) or high light (HL, 300 µmol m^−2^ s^−1^) for 2 days in the growth chamber. Anthocyanin phenotype of seedlings was examined and photographed under a dissection microscope. Plants were grown in soil under light intensity of 100 µmol m^−2^ s^−1^ in long-day conditions (16 h light / 8 h dark) at 22°C in a phytotron for seeds harvesting.

### Anthocyanin content measurement

Anthocyanin content was measured using a previously described method (Isaac and Alberto, 1986). Seedlings weighed about 20 mg were frozen in liquid nitrogen and ground into powder, then added 750 µL of 0.1% (v/v) HCl methanol solution. Samples were placed in the dark overnight. After adding 500 µL distilled water and 500 µL chloroform, and centrifugation at 12000 g for 4 min at room temperature, the absorbance of the supernatants were measured at 530 and 657 nm by a spectrophotometer. (A_530_ - 0.25 × A_657_) per gram fresh weight was used to quantify the relative amounts of anthocyanins.

### Chlorophyll content measurements

Total chlorophyll levels were extracted by homogenizing the seedlings in 95% (v/v) ethanol, and leaving the solutions at 4 °C for overnight incubation. The absorbance was then detected at wavelengths OD_665_ and OD_649_ with a spectrophotometer (Manufacture). Chlorophyll a (µg mL^-1^) was calculated by (13.95 × OD_665_) — (6.88 × OD_649_) (mg/L), Chlorophyll b (µg mL^-1^) was calculated by (24.96 × OD_649_) — (7.32 × OD_665_). Then converted according to the fresh weight of the material.

### H_2_O_2_ content quantification

Seedlings weighed about 50 mg were frozen in liquid nitrogen and ground into powder and homogenized in 1 ml of cold acetone. Then, H_2_O_2_ content was determined using a hydrogen peroxide assay kit (Solarbio, BC3590) and its absorbance was measured at 415 nm. Data are represented as the amount of H_2_O_2_ per gram leaf (µmol/g).

### Gene expression analyses

RNA samples were extracted with RNAiso Plus (Takara) and cDNA was synthesized from 1000 ng of total RNA using a Hifair^®^ III 1st Strand cDNA Synthesis SuperMix (gDNA digester plus) (Yeasen, 11141ES60). Quantitative RT-PCR was performed with cDNA templates, gene-specific primers, and Hieff UNICON^®^ Universal Blue qPCR SYBR Master Mix (Yeasen, 11184ES08) in a total volume of 20 uL. *ACTIN7* was used as an internal control for most analyses. For samples treated with HL, *CYCLOPHILIN 5* (*CYP5*) was used as internal control, as its expression was shown to be unchanged in HL-exposed leaves(Galvez-Valdivieso et al., 2009). The primers used are listed in **Supplemental Table 1**. Statistical *t* test analyses were calculated using Excel and ANOVA analyses were performed using SPSS. Data with *P* value lower than 0.05 was considered to be statistically significant. The results of statistical analyses are shown in **Supplemental Table 2**.

### Electrophoretic mobility shift assay (EMSA)

Probes were PCR amplified using Cy5-labeled forward primer pairs, from synthetic complementary oligonucleotides of *DFR*, LDOX, UF3GT and *TTG1* (or mDFR with GLK2 binding motif mutation). For protein, the sequence encoding GLK2 was cloned into the pCold-TF vector, expressed in *E. coli*, and purified with Ni-NTA His•Bind^®^ Resin agarose (Merck, 70666). The binding reaction was performed in 20 µL of binding buffer [10 mM Tris (pH 8.0), 1 mM KCl, 4 mM MgCl_2_, 0.5 mM DTT, 5% (v/v) glycerol, 0.2 mM EDTA, and 0.01% BSA], using 5 nM probe and 200 ng protein, and incubated at room temperature (25°C) for 30 min. The Cy5-labeled DNA after binding reactions were resolved by electrophoresis in 5% (v/v) native polyacrylamide gel at 4°C, and the gel was scanned by a Amersham Typhoon 5 Biomolecular Imager.

### Chromatin immunoprecipitation (ChIP)

The ChIP experiment was carried out as described previously(Hu & Xu, 2016). For ChIP assays, the surface-sterilized *glk1 glk2* double mutant and double mutant lines overexpressing GLK2 (35S:GLK2) seeds were treated at 4°C in the dark for 3 days, and grown on half-strength Murashige and Skoog (MS) medium (pH 5.8) containing 0.8% agar and 1% sucrose at 22°C under light intensity of 100 µmol m^−2^ s^−1^ in long-day conditions (16 h light / 8 h dark) for 2 weeks in the growth chamber. Four grams of seedling tissue from *glk1 glk2* and 35S:GLK2 plants were harvested and crosslinked with 1% (v/v) formaldehyde (Sigma-Aldrich) for 20 min under a vacuum. Cross-linking was stopped by adding Gly to afinal concentration of 0.125 M. The seedlings were rinsed with water, frozen in liquid nitrogen, and ground to a fine powder. The powder was homogenized in nuclear extraction CLB buffer (50 mM HEPES [pH 8.0], 1mM EDTA, 150mM NaCl, 10% (v/v) Glycerol, 1% (v/v) TritonX-100, and 0.1 mM protease inhibitor cocktail tablets [Roche]). Nuclei were precipitated by centrifugation at 3000g for 20 min at 4°C, washed with nuclear extraction buffer, and lysed in nuclei lysis buffer (CLB buffer and 1% [w/v] SDS). The chromatin was sheared by sonication to around 500 bp. The chromatin solution was diluted 10- fold with CLB buffer. Anti-GLK2 antibody prebound to protein A/G sepharose was mixed with the chromatin solution and incubated overnight at 4 °C. Immunocomplexes were precipitated and washed with four different buffers: low-salt buffer (50 mM HEPES [pH 7.5], 150 mM NaCl, 1 mM EDTA), high-salt buffer (50 mM HEPES [pH 7.5], 500 mM NaCl, 1 mM EDTA), LiCl washing buffer (10 mM Tris-HCl [pH 8.0], 0.25 M LiCl, 0.5% [v/v] Nonidet P-40, 1 mM EDTA), and transposable element washing buffer (10 mM Tris-HCl [pH 8.0], 1 mM EDTA). The bound chromatin fragments were eluted with elution buffer (100 mM NaHCO_3_, 1% [w/v] SDS), and the cross-linking was reversed by incubating overnight in a final concentration of 200 mM NaCl at 65 °C. The mixture was treated with Proteinase-K to remove proteins. The genomic DNA was purified with an equal volume of phenol/chloroform/isoamyl alcohol and precipitated with 80mM NaAc. of 100% ethanol at -20 °C for 2 h to overnight. The sample was centrifuged at 16,000g for 12 min at 4 °C to recover the DNA, which was dissolved in 20 ul of double distilled water. About 10% of sonicated but non-immunoprecipitated chromatin was reverse cross linked for using as an input DNA control. Both immunoprecipitated DNA and input DNA were analyzed by Quantitative RT-PCR using primers for target gene promoter regions. The enrichment of GLK2 binding promoter regions were evaluated by the ratio between the immunoprecipitated and input reactions. The primer pairs used in the ChIP experiments are listed in Supplemental Table 1.

### Transient transcription dual-luciferase assay

The protoplast isolation and PEG-mediated transformation were carried out as previously described(Xu Li et al., 2020). We isolated protoplasts from two-week-old Col-0 plants grown on half-strength Murashige and Skoog (MS) medium (pH 5.8) containing 0.8% agar and 1% sucrose at 22°C under light intensity of 100 µmol m^−2^ s^−1^ in long-day conditions (16 h light / 8 h dark) for 2 weeks in the growth chamber. The promoter region of *DFR*, *LDOX*, *UF3GT*, *PAP1*, *TT8*, and *TTG1* were placed into the pGreenII 0800-LUC vector (Firefly luciferase) for reporter constructs; the full length CDS of GLK2 was placed into the pGreenII 62SK-35S:: vector for effector construct. Protoplasts were transfected with a total of 20 μg DNA of reporter constructs with effector construct (empty effector construct was used as internal control) and incubated overnight. Then the reporter and effector proteins were released from the protoplasts, and the ratio of Firefly luciferase to Renilla luciferase was measured and calculated by using a luminometer (GloMax 20/20, Promega), with a Dual-Luciferase Reporter Assay System (E1910, Promega), according to the manufacturer’s instructions.

### Protein pull-down assay

Full-length coding sequences of *GLK1* and *GLK2* were cloned into the pGEX-6P-1 vector, and full-length coding sequence of *TT8* was cloned into the pCold-TF vector. GST-GLK1, GST-GLK2, and His-TT8 proteins were expressed in *E.coli* strain *BL21*. The *E.coli BL21* containing empty pGEX-6P-1 vector was used as negative control (producing N- terminal GST protein only). Next, 0.5 ml of each *E.coli* strain (one GST tagged plus one His tagged) was collected, mixed, and re-suspended in PBS buffer [1.4M NaCl, 2.7mM KCl, 100mM Na_2_HPO_4_, 18 mM KH_2_PO_4_ (pH7.3), 1mM DTT, 0.2% Triton X-100, and 1% cocktail protease inhibitor (Roche)]. The cells were broken with a sonicator for 3 times (30s on and 60s off) on ice, the cellular debris was removed by centrifugation at 18000 g for 30 min, then 10 μL of Glutathione Sepharose 4B beads (17-0756-01, GE Healthcare) was added to the mixture sample and rotated at 4°C for 16 hours. Unbound proteins were removed by washing the beads with PBST buffer [100 mM Tris (pH8.0), 5 mM EDTA, 10 mM glutathione, and 0.1% Triton X-100] for at least 4 times. Retained proteins were released by boiling in loading buffer at 95°C for 5 min, then loaded on SDS-PAGE for Western blot. The anti-GST antibody (Santa Cruz Biotechnology) was used to detect GLK1 and GLK2 proteins that fused with a N-terminal GST tag, and the anti-His antibody (Abmart) was used to detect whether TT8 protein was pulled down by GST tagged GLK proteins.

### Bi-luminescence complementation (BiLC) assay

GLK2 or TT8 was fused to the N- or C-terminus of Firefly luciferase, and the constructs were transformed into *Agrobacterium tumefaciens* strain *GV3101*. Overnight cultures of *Agrobacteria GV3101* were collected by centrifugation at 4000 g for 10 min, and resuspended in MES buffer (10 mM MES,10 mM MgCl_2_, and 100 mM acetosyringone). They were mixed with *Agrobacteria* expressing pSoup-P19 to a final OD600=0.5, and incubated at room temperature for 3 h in the dark before infiltration. The *Agrobacterium* suspension in a 1 ml syringe (without the metal needle) was carefully press-infiltrated into healthy leaves of 3-week-old tobacco *N. benthamiana*. The infiltrated plants were returned to long-day conditions for 3 days. Luciferase activity can be imaged with a CCD camera (Tanon-5200, BioTanon), 10 min after leaves were infiltrated with luciferin solution.

### Accession numbers

Sequence data for genes described in this article can be found in the Arabidopsis Information Resource under the following accession numbers: *GLK1* (AT2G20570), *GLK2* (AT5G44190), *DFR* (AT5G42800), *LDOX* (AT4G22880), *UF3GT* (AT5G54060), *PAP1* (AT1G56650), *TT8* (AT4G09820), *GL3* (AT5G41315), *EGL3* (AT1G63650), *TTG1* (AT5G24520), *ACTIN7* (AT5G09810), *CYCLOPHILIN 5* (AT2G29960).

## Supplemental data

Supplemental Figure 1. GLKs positively regulate anthocyanin accumulation during the first day of seedling de-etiolation in Arabidopsis

Supplemental Figure 2. Light regulation of AtGLK1 and AtGLK2 genes

Supplemental Figure 3. Loss of GLK2 exceptionally delays the deetiolation process of Arabidopsis seedlings

Supplemental Figure 4. GLK2 positively regulates MBW complex member genes

Supplemental Figure 5. GLK2 actives the expression of DFR gene independent from PAP1 and TTG1 of the MBW complex

Supplemental Figure 6. GLK2 positively regulates anthocyanin accumulation in mature Arabidopsis

Supplemental Figure 7. Detection of GLK2 protein by anti-GLK2 antibodies

Supplemental Table S1. Primer used in this study.

Supplemental Table S2. The results of statistical analyses.

## Author contributions

XZ and PW designed the study. LY performed ChIP and RZ performed q-PCR experiments. XZ worked on all the experiments. XZ and PW analyzed the data and wrote the manuscript.

## Acknowledgments

We thank Prof. Xinguang Zhu for fruitful discussions on the work. This work was supported by the National Natural Science Foundation of China (No. 31970257).

---

**Supplemental Figure 1.**
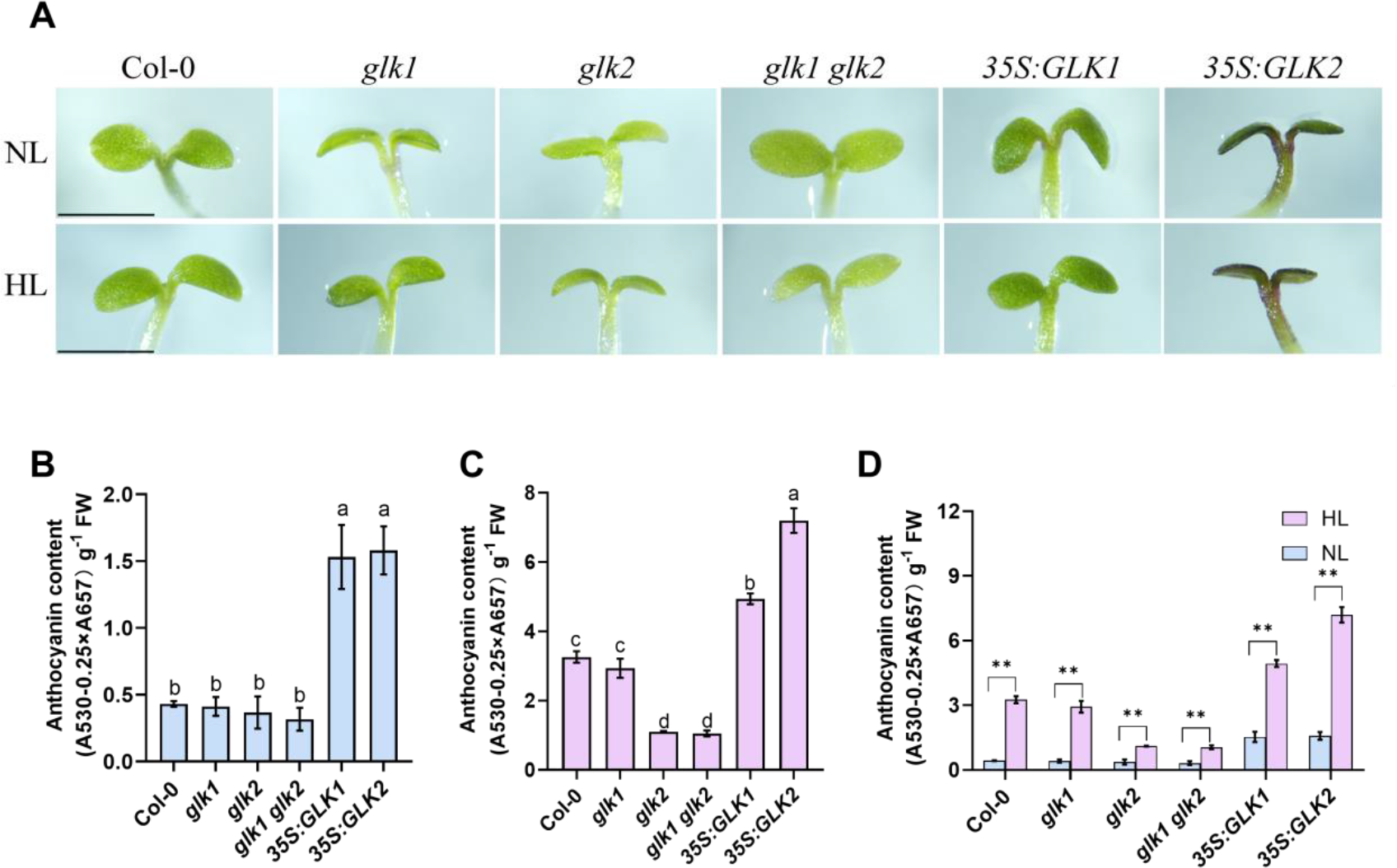
GLKs positively regulate anthocyanin accumulation during the first day of seedling de-etiolation in Arabidopsis **(A)** Phenotypes of wild-type (Col-0), *glk1*, *glk2*, double mutant (*glk1 glk2*), and double mutant lines overexpressing either *GLK1*(*35S:GLK1*) or *GLK2* (*35S:GLK2*). Seedlings germinated in darkness for 4 d were transferred to normal light (NL, 100 μmol m^-2^ s^-1^) and high light (HL, 300 μmol m^-2^ s^-1^) for 1 d in the growth chamber. Experiments were repeated three times and similar results were obtained. Scale represents 1 mm. **(B)** The anthocyanin content of NL grown seedlings were analyzed. **(C)** The anthocyanin content of HL grown seedlings were analyzed. **(D)** Comparison of the NL and HL effect on anthocyanin content. Error bars represent mean ± SD (n=3). Statistical analysis in (B) and (C) was performed using ANOVA with Turkey’s HSD test; *P* < 0.05, different letters indicate statistically significant difference. Asterisks in (D) indicate significant differences with \*\**P* < 0.01 (Student’s *t*-test).

**Supplemental Figure 2.**
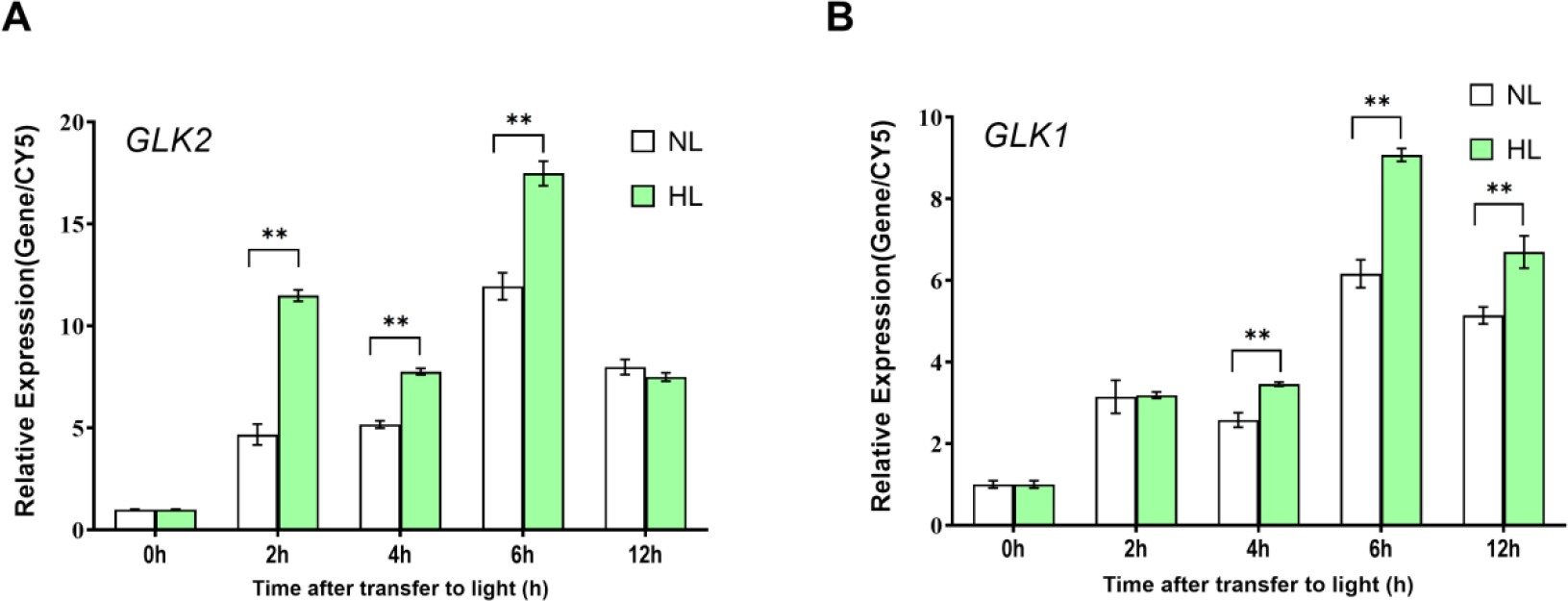
L**i**ght **regulation of *AtGLK1* and *AtGLK2* genes.** Relative expression of *GLK2* **(A)** and *GLK1* **(B)** genes after 4 d dark-grown wild-type seedlings were transfer to normal light (NL, 100 μmol m^-2^ s^-1^) or high light (HL, 300 μmol m^-2^ s^-1^) for 0, 2, 4, 6, and 12 h. The CY5 gene was analyzed as an internal control, and the expression levels at 0 h were set to 1. Error bars represent mean± SE (n = 3). Asterisks indicate a significant differences (*P<0.05, **P<0.01, Student’s *t*-test).

**Supplemental Figure 3.**
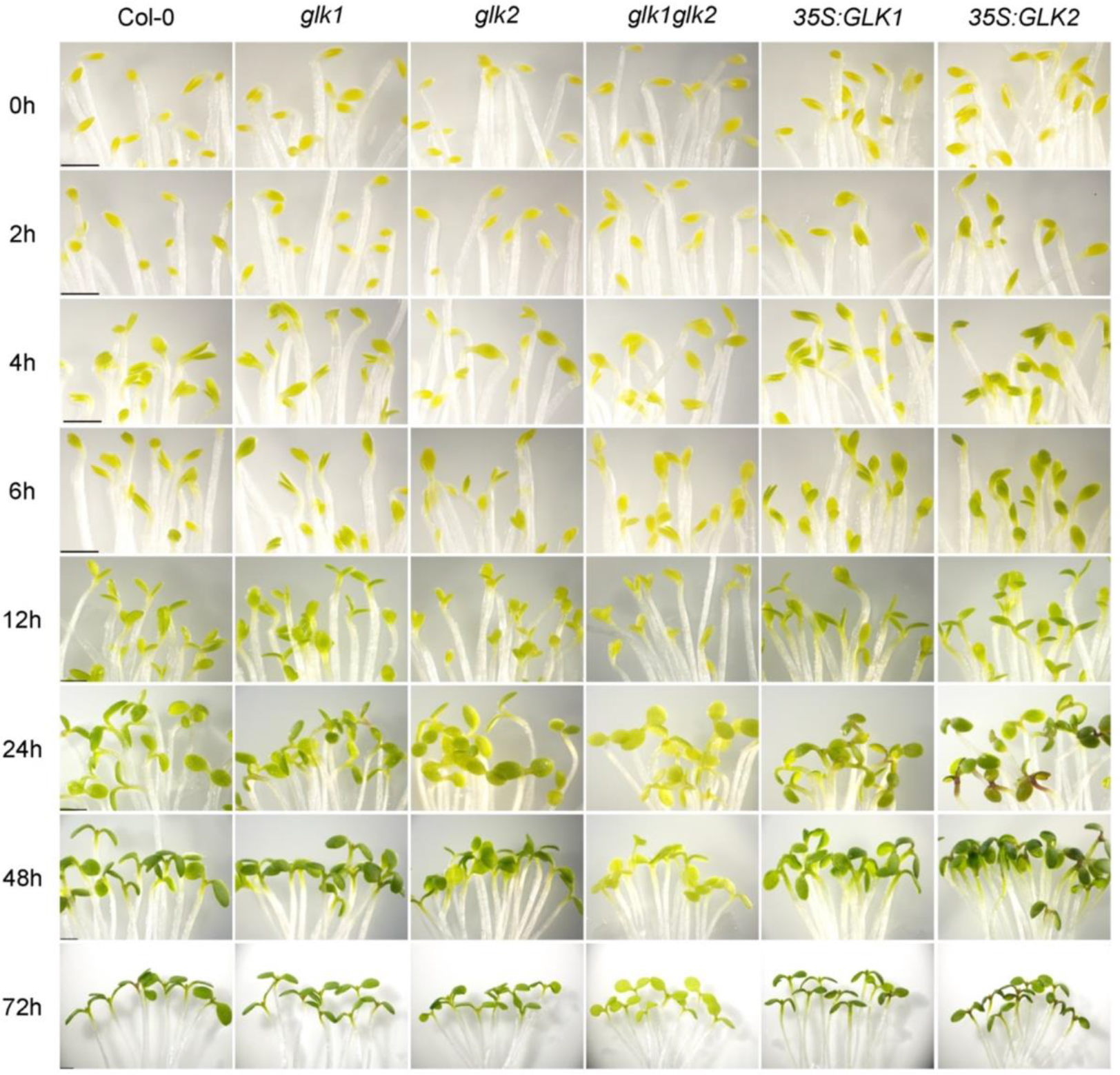
Loss of *GLK2* exceptionally delays the deetiolation process of Arabidopsis seedlings. 4-day-old dark-grown seedlings were transferred to 100 μmol m^-2^ s^-1^ white light for the times indicated. Scale bar represents 1 mm. Over 15 seedlings were analyzed for each genotype. Experiments were repeated five times and similar results were obtained.

**Supplemental Figure 4.**
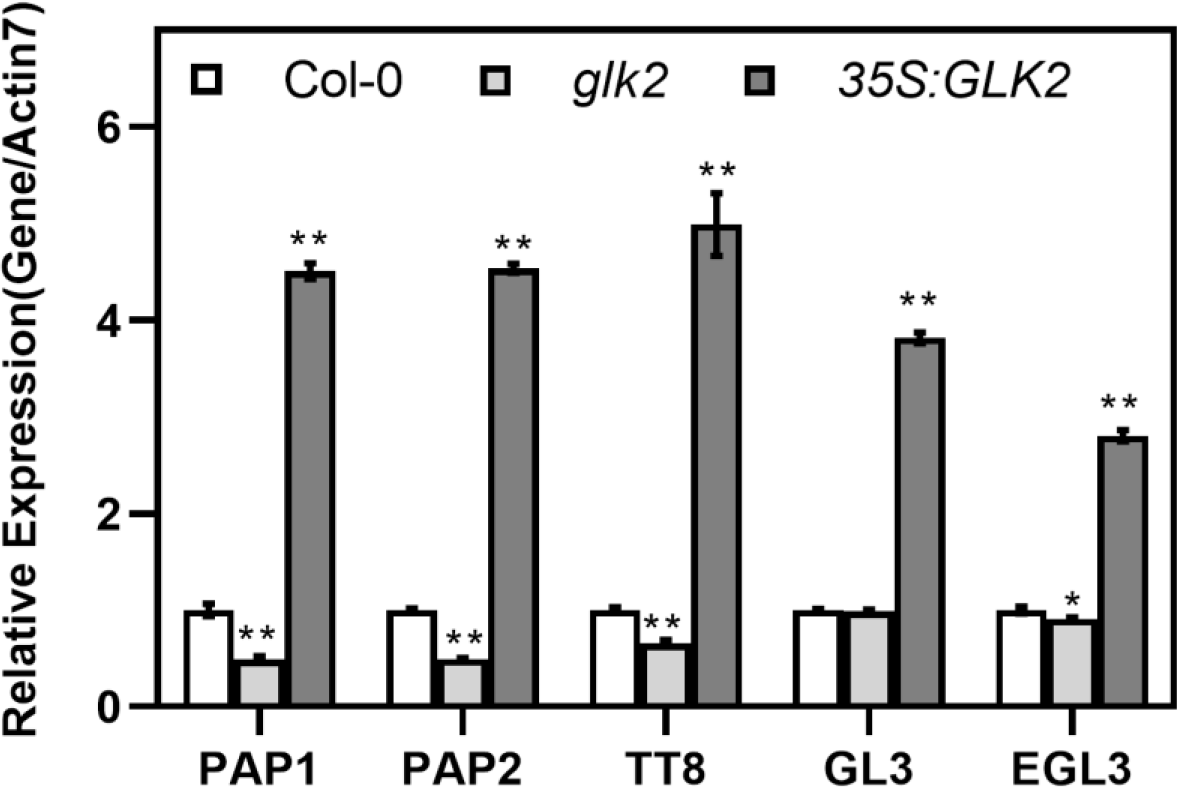
GLK2 positively regulates MBW complex member genes. Transcript levels of MBW complex genes were tested using RT-qPCR for seedlings of Col-0, *glk2*, and *35S:GLK2* grown under high light (HL, 300 μmol m^-2^ s^-1^) for 2 days after germination in darkness for 4 d. The Actin7 gene was analyzed as an internal control, and the expression levels of Col-0 were set to 1. Error bars represent mean±SE (n = 3). Asterisks indicate a significant differences (*P<0.05, **P<0.01, Student’s *t*-test).

**Supplemental Figure 5.**
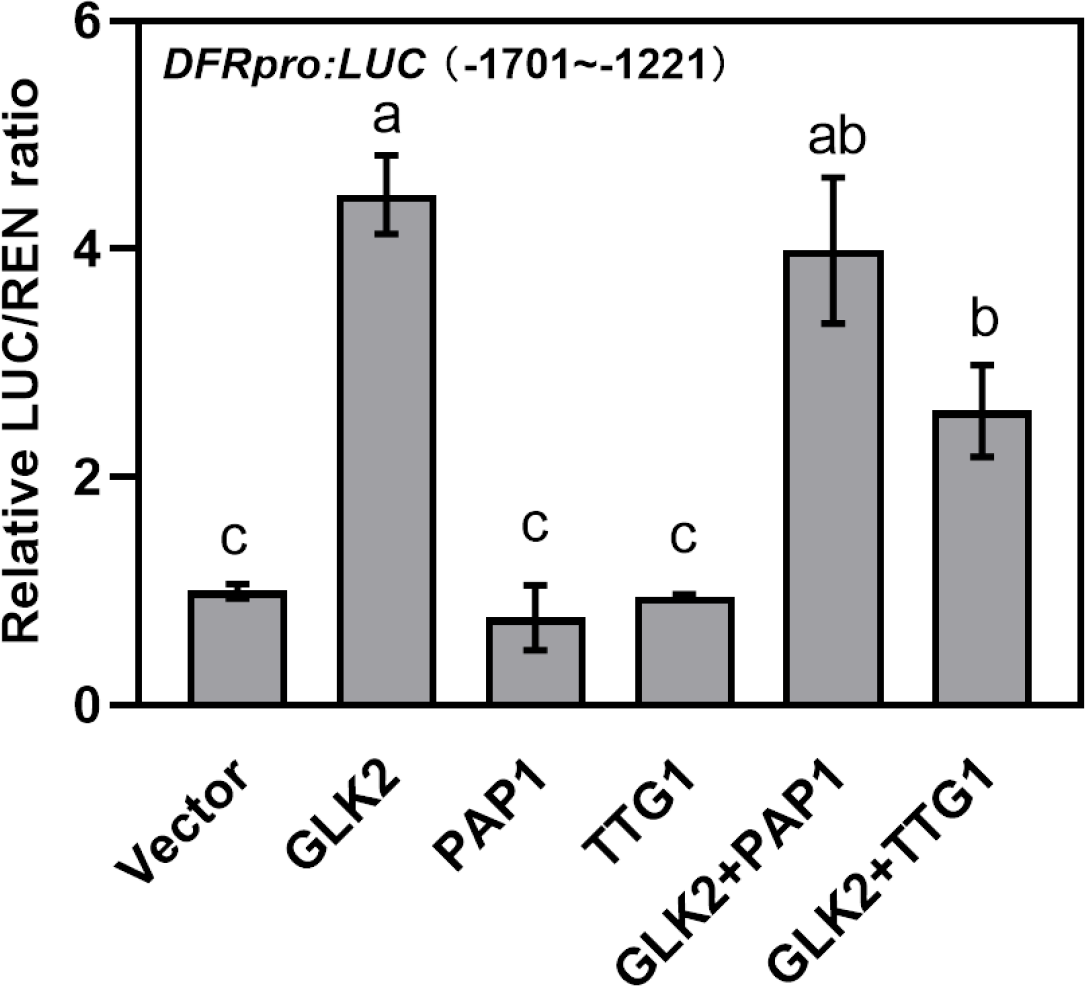
GLK2 actives the expression of *DFR* gene independent from PAP1 and TTG1 of the MBW complex. Dual-luciferase assays were performed similar as in Figure 5D. Error bars represent mean±SD (n=3). Letters “a” to “c” indicate statistically significant difference, as determined by one-way ANOVA, followed by Tukey’s HSD test (P < 0.05).

**Supplemental Figure 6.**
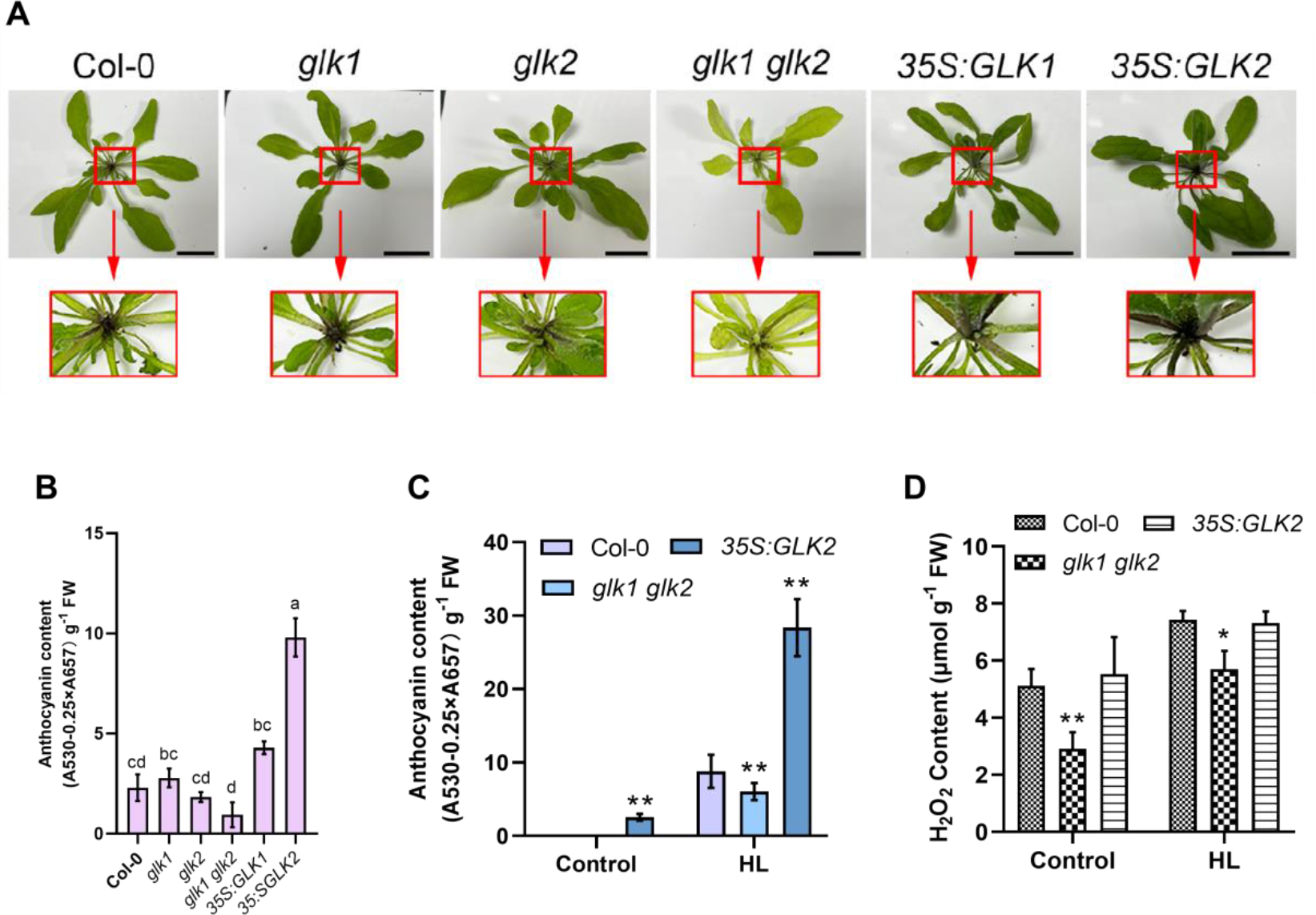
GLK2 positively regulates anthocyanin accumulation in mature Arabidopsis. **(A)** Anthocyanin accumulation in leaf petioles of 5-week-old WT, *glk1*, *glk2*, double mutant (*glk1 glk2*), and double mutant lines overexpressing either *GLK1*(*35S:GLK1*) or *GLK2* (*35S:GLK2*). The red squares show the magnified images. Scale bar represents 2 cm. **(B)** Quantified anthocyanin accumulation in leaf petioles of 5-week-old Arabidopsis plants. **(C)** Five-week-old Col-0, *glk2*, and 35S:GLK2 plants grown under light intensity of 100 μmol m^-2^ s^-1^ long-day conditions (16 h light/8 h dark) were transferred to high light (HL, 500 μmol m^-2^s^-1^) for 2 d. The anthocyanin contents were analyzed. **(D)** H_2_O_2_ quantification of the plants in (C). Error bars represent mean±SD (n=3). Statistical analysis in (B) was performed using ANOVA with Turkey’s HSD test; *P* < 0.05, different letters indicate statistically significant difference. Asterisks in (C) and (D) indicate significant differences with \**P* < 0.05 and \*\**P* < 0.01 (Student’s *t*-test), respectively.

**Supplemental Figure 7.**
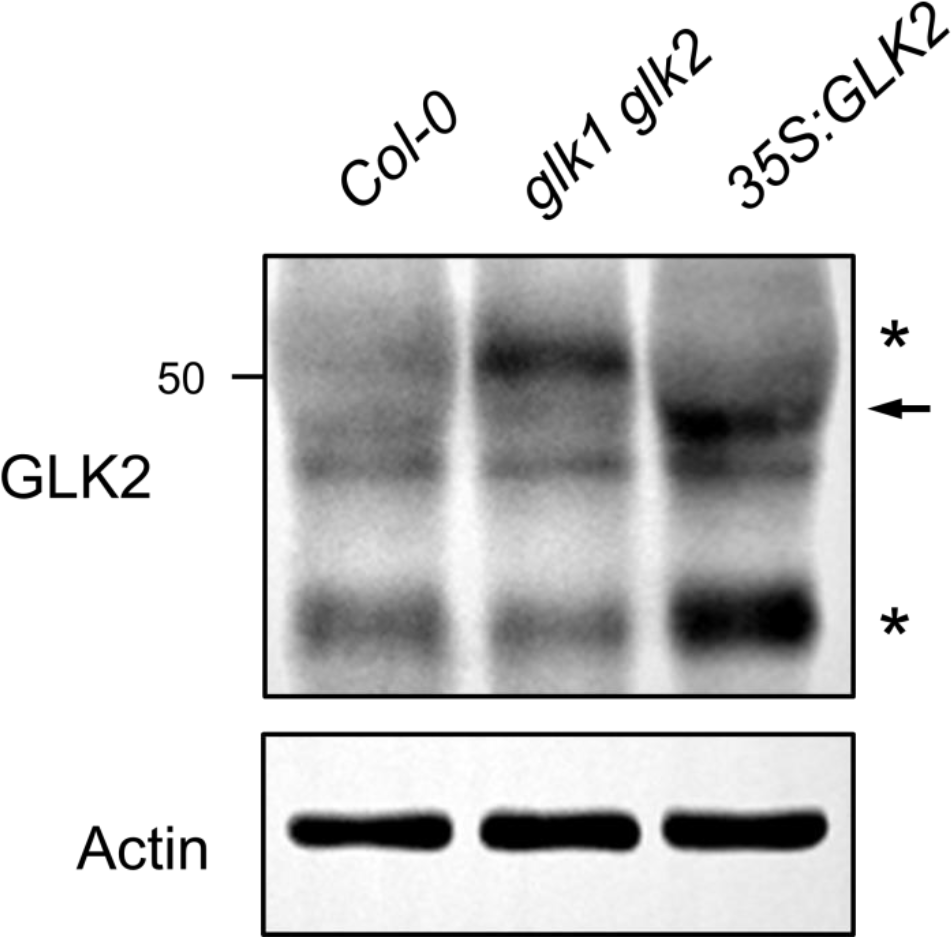
Detection of GLK2 protein by anti-GLK2 antibodies. Total protein were extracted from 2-week-old Arabidopsis seedlings wild-type (Col-0), double mutant (*glk1 glk2*), and double mutant lines overexpressing *GLK2* (*35S:GLK2*) grown in growth chamber, resolved by SDS-PAGE, and probed with antibodies against GLK2. The arrow indicates GLK2 proteins. Asterisks indicate nonspecific proteins detected by antibodies.

